# Co-estimation of Phylogeny-aware Alignment and Phylogenetic Tree

**DOI:** 10.1101/077503

**Authors:** Chunxiang Li, Alan Medlar, Ari Löytynoja

## Abstract

The phylogeny-aware alignment algorithm implemented in both PRANK and PAGAN has been found to produce highly accurate alignments for comparative sequence analysis. However, the algorithm’s reliance on a guide tree during the alignment process can bias the resulting alignment rendering it unsuitable for phylogenetic inference. To overcome these issues, we have developed a new tool, Canopy, for parallelized iterative search of optimal alignment. Using Canopy, we studied the impact of the guide tree as well as the number and relative divergence of sequences on the accuracy of the alignment and inferred phylogeny. We find that PAGAN is the more robust of the two phylogeny-aware alignment methods to errors in the guide tree, but Canopy largely resolves the guide tree-related biases in PRANK. We demonstrate that, for all experimental settings tested, Canopy produces the most accurate sequence alignments and, further, that the inferred phylogenetic trees are of comparable accuracy to those obtained with the leading alternative method, SATé. Our analyses also show that, unlike traditional alignment algorithms, the phylogeny-aware algorithm effectively uses the information from denser sequence sampling and produces more accurate alignments when additional closely-related sequences are included. All methods are available for download at http://wasabiapp.org/software.

## 1 Introduction

The phylogeny-aware alignment algorithm implemented in PRANK (Löytynoja and Goldman 2005, 2008) has been shown to produce highly accurate alignments for comparative analyses of gene sequences (e.g., Fletcher and Yang 2010; Markova-Raina and Petrov 2011; Jordan and Goldman 2012; Privman et al. 2012). Such comparative studies are typically performed on species whose phylogeny is known *a priori* and alignments can be inferred using a trusted phylogeny as the alignment guide tree. This is in contrast to phylogenetic inference where the tree is the parameter of interest and therefore not known beforehand. For the phylogeny-aware algorithm, this circularity poses a challenge as the guide tree necessarily biases the alignment, the accuracy of which, will impact the inferred phylogeny.

Progressive alignment algorithms build a multiple sequence alignment from a set of pairwise alignments performed according to a binary guide tree (Hogeweg and Hesper 1984). Starting from the leaves of the guide tree, each alignment coalesces two evolutionary lineages and creates a new node that itself can be merged with another lineage (Löytynoja and Goldman 2009). Most implementations of progressive alignment consider the guide tree as a nuisance parameter that should be hidden from the user and represent the internal nodes of the alignment with sequence profiles (e.g., Edgar 2004; Larkin et al. 2007; Katoh and Standley 2013). From an evolutionary point of view, the internal nodes are not averages of their descendants, but ancestors whose states can be inferred.

Phylogeny-aware alignment programs, such as PRANK, aim to accurately reconstruct the ancestral sequence history. While the reconstruction of ancestral character states is relatively straightforward, the inference of insertions and deletions necessitates the use of outgroup information from the subsequent alignment to ensure gap placement is evolutionarily consistent (Löytynoja and Goldman 2005). When sequences are densely sampled and the chances of near-by mutation events are low, this approach provides an accurate inference of ancestral sequences. However, errors in the guide tree make outgroup information misleading, causing errors in reconstructed ancestors and affect subsequent alignment steps (Löytynoja 2014).

In relation to phylogenetic analysis, the circularity of requiring a guide tree to create a multiple sequence alignment prior to phylogenetic inference is a serious, but often overlooked, issue. While guide tree bias affects all progressive alignment algorithms, PRANK is acutely affected. Another phylogeny-aware alignment method, PAGAN (Löytynoja et al. 2012), is a recent attempt at ameliorating the issues related to guide tree bias. Unlike PRANK, which makes decisions based on local information only, PAGAN defers the assignment of alignment gaps as insertions or deletions until sufficient evidence is available from throughout the alignment. However, there are currently no published results evaluating PAGAN’s performance in the context of phylogenetic inference.

Recent work by Liu *et al*. showed that the error in phylogenetic analysis can be reduced by performing successive alignment and phylogenetic inference steps, with the phylogenetic tree generated in one iteration used as the alignment guide tree in the next (Liu et al. 2009, 2012). Their tool, SATé, scales to large data sets using a divide-and-conquer strategy to partition the dataset into subsets, aligns them separately and then merges the sub-alignments into a complete multiple sequence alignment. While SATé also incorporates a version of PRANK for alignment, it merges the the results using a traditional profile alignment method and the resulting full alignments are not comparable to standard PRANK alignments.

Given the details of the phylogeny-aware alignment algorithm, and its misuse in SATé, it is not clear whether PRANK’s dependence on a guide tree can be overcome, while retaining alignment accuracy, and enable its use in phylogenetic analysis. To investigate further, we implemented the necessary features needed to perform an iterative SATé-like divide-and-conquer procedure using PRANK in a new tool called Canopy. Canopy is capable of generating larger multiple sequence alignments than PRANK due to distributing the workload across multiple CPU cores. In the following sections we first briefly describe the functionality of Canopy and use simulated data to assess the performance of all phylogeny-aware methods for phylogenetic analysis.

## 2 Description and Benchmark

### 2.1 Canopy: Parallelized, Iterative Search of Alignment and Phylogeny

Canopy utilizes existing analysis tools to infer and optimize a multiple sequence alignment and phylogenetic tree. The two main functions of Canopy are: (i) to break up the phylogeny-aware alignment problem into a set of smaller subtasks and merge the resulting solutions into a full alignment and (ii) to iterate between the alignment and phylogenetic inference steps and output the highest scoring solution. Canopy allows users to run one or both functions independently to accommodate different use cases to, for example, parallelize a large PRANK alignment without inferring a phylogenetic tree. In other circumstances, if the alignment program natively supports multithreading, like PAGAN for example, or the data set is small enough; Canopy need only iterate between alignment and phylogenetic inference.

The main difference between Canopy and SATé is the use of phylogeny in the alignment. SATé uses the guide phylogeny only to define the sequence subsets and their relationships, but then ignores the original phylogeny during each subalignment. In contrast, Canopy uses the guide phylogeny together with each sequence subset to guide their alignment. Afterwards, Canopy merges the subalignments, respecting their internal phylogenies as well as relative phylogenetic relationships. To accomplish this, we implemented functions in PRANK to merge two intermediate results enabling the phylogeny-aware alignment algorithm to proceed as usual. Indeed, Canopy outputs identical results to PRANK, but is faster due to using multiple CPUs. A detailed description of the new methods in both Canopy and PRANK is given in Supplemental Material.

Though Canopy supports parallelism, numerous factors influence the asymptotic performance behavior as we utilize increasing numbers of CPUs (Fig. 1). Firstly, the number of independent alignments that can be performed concurrently is constrained by the guide tree topology. If the guide tree is unbalanced, then there are fewer pairwise alignments (and therefore sequence subsets) that can be aligned independently. Indeed, in the pathological case of a perfectly unbalanced comb-like topology, only a single CPU can be used because all pairwise alignments would need to be performed in sequential order. Secondly, the merging procedure is limited to two alignments at a time and the repeated reconstruction of sequence history adds substantial overhead. In this sense, Canopy must tradeoff between using smaller sequence subsets to increase parallelism and larger subsets to reduce the fixed overhead from merge operations.

**Figure 1:**
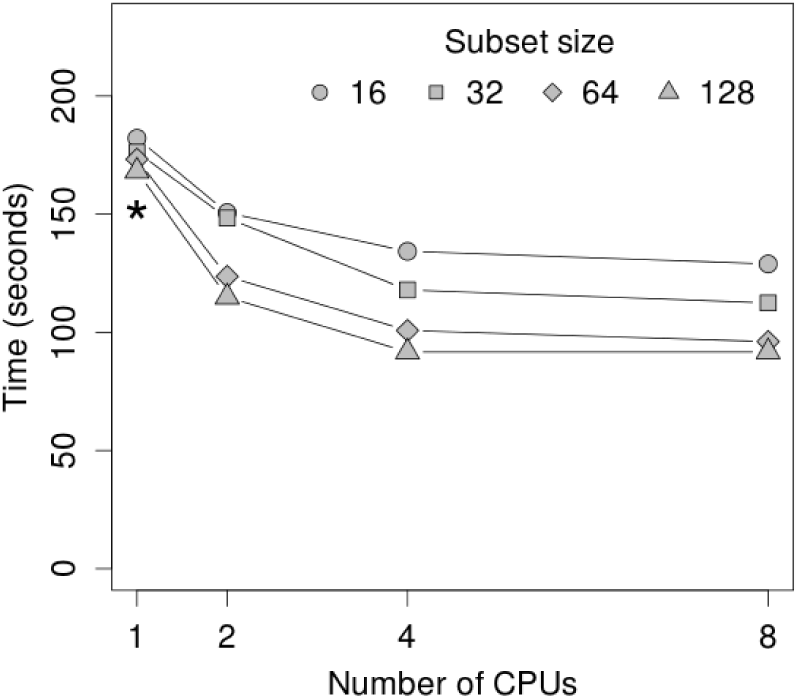
Time (y axis; seconds) taken by Canopy to align 512 sequences using different numbers of CPU cores (x axis) and sizes of sequence subsets (different symbols). The asterisk indicates the original PRANK. The guide tree is perfectly balanced.

### 2.2 Accuracy of Inferred Alignment and Phylogeny

We assessed the performance of iterative alignment with Canopy using simulated sequence data, focusing on the accuracy of the inferred alignments and phylogenetic trees. We used INDELible ver.1.03 (Fletcher and Yang 2009) to simulate DNA data sets using the HKY substitution model (kappa = 2.5, base frequencies = 0.29, 0.21, 0.29 and 0.21 for T, C, A and G, respectively). The indel rate was set to 0.05 and the length of indels to follow the power-law distribution with a shape parameter of 1.7 and maximum length of 50. Starting from a root sequence of 500 nucleotides, we simulated data sets of 800 sequences. The sequence data were simulated using ten randomly generated rooted trees. For each of the ten trees, the tree branches were rescaled to make the maximum root-to-tip distance be 0.5, 1.0 and 1.5, which we refer to as ‘Close’, ‘Intermediate’ and ‘Distant’, respectively. Ten replicate alignments were generated for each rescaled tree giving 100 distinct alignments of 800 sequences at three different evolutionary divergence levels. The simulated data sets were then down-sampled by repeatedly keeping every second sequence, producing additional data sets of 400, 200, 100 and 50 sequences.

We assessed the alignment accuracy using MetAl ver.1.1.0 (Blackburne and Whelan 2012) and measured the correctness of phylogenetic trees, inferred using RAxML ver.7.3.5 (Stamatakis 2014), using the Robinson-Foulds metric (Robinson and Foulds 1981) as implemented in the DendroPy package ver.3.12.0 (Sukumaran and Holder 2010). We normalized the Robinson-Foulds metric by dividing by 2(*n−*3), where *n* is the number of taxa in the tree, giving the percentage of false or missing splits in the inferred tree in comparison to the true tree. We assessed alignment and topological accuracy on the 50 taxa that were shared between all data sets. We argue that this provides a more appropriate measure of accuracy compared to analyzing the complete results because, as we simulate data sets with a set root-to-tip distance, the accuracy measurements for larger (and therefore denser) data sets get “crowded out” by the easy cases of more similar sequences. Furthermore, by focusing on the 50 shared taxa only, we effectively assess the accuracy of the major evolutionary lineages that the users most care about.

We compared the performance of Canopy to that of standard PRANK ver.130807 and PAGAN ver.0.47, and of SATé ver.2.0 (Liu et al. 2012). With PRANK and PAGAN, we performed the alignments using a guide tree inferred from the data (both programs internally use neighbor joining (NJ) (Saitou and Nei 1987)). As Canopy is focused on improving the performance of PRANK, its starting tree is the same as PRANK’s default NJ tree. To provide a baseline for Canopy, we additionally ran PRANK with the true simulation tree as the guide tree (from now on simply referred to as the “true tree”), reflecting performance given ideal circumstances. PRANK, PAGAN and SATé were run with default parameter settings. Both iterative methods, Canopy and SATé, were run for 25 iteration and the maximum likelihood (ML) score from RAxML was used as the optimization criteria. The assumption is that the ML score is a good surrogate for the accuracy of alignment and the tree, the two features that are of interest but which cannot be measured in real-life analyses.

Our first finding is that the accuracy of phylogeny-aware sequence alignment improves with more densely sampled input data (Fig. 2). This trend is expected in phylogenetic inference (Zwickl and Hillis 2002), but not necessarily with all progressive alignment methods. For the set of 50 sequences shared by data sets of different sizes, median alignment error for PRANK using the true tree decreases with the number of sequences in the input set.

**Figure 2:**
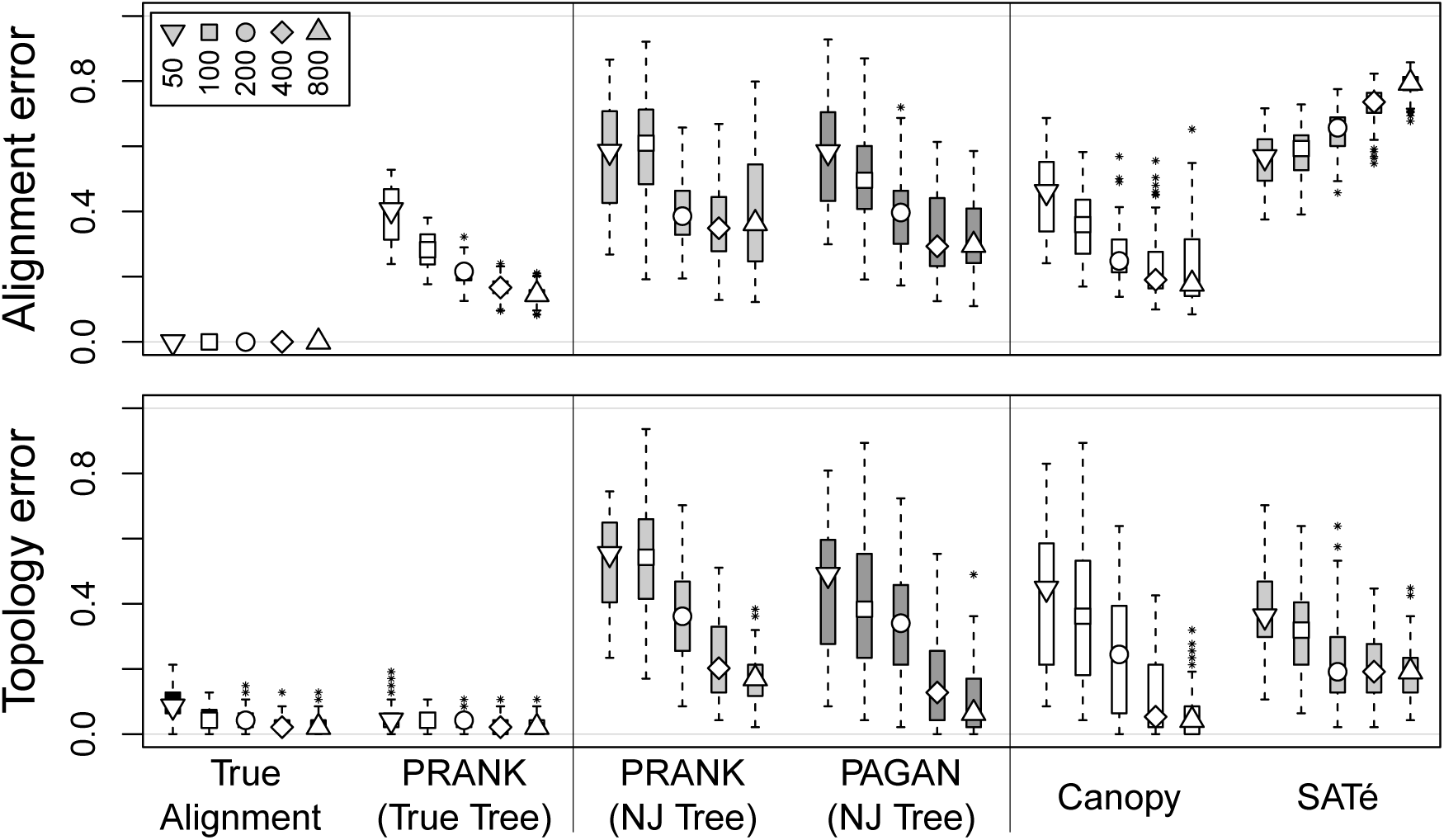
Alignment and topology error for ‘Intermediate’ data sets. The five symbols from left to right refer to data sets of 50, 100, 200, 400 and 800 sequences. The symbols indicate the median values and the boxes show the second and fourth quartiles. The method names are given at the bottom.

The true tree is unknown in most real-life analyses. Despite this fact, however, using a guide tree inferred from the data, the trend of alignment error decreasing as sequence sampling density increases is observed for all phylogeny-aware methods (Fig. 2). Compared to the other phylogeny-aware alignment programs, Canopy consistently outperforms both standard PRANK and PAGAN in terms of median alignment accuracy. In contrast to this, SATé alignment error actually *increases* with the number of input sequences. These trends are replicated across all three data sets, irrespective of simulated substitution rate (see Supplemental Material).

The relative performance of the different approaches in terms of topological accuracy is often contrary to alignment accuracy. For example, while SATé produces more erroneous alignments than standard PRANK (with NJ tree) for Intermediate and Close data sets, these alignments produce more accurate phylogenetic trees (Fig. 2, Fig. S1). This difference can be attributed to errors in the guide tree biasing the alignment. When the true tree is used, phylogenies inferred from PRANK alignments can be more correct than even those inferred from true alignments (Fig. 2). In contrast, the error in SATé alignments is expected to be more random and, therefore, does not bias the inferred phylogenies strongly in any particular direction. Canopy appears to overcome some of this bias and produces highly similar or more accurate phylogenetic trees compared to SATé for higher numbers of sequences (400 and 800) across all substitution rates. For lower numbers of sequences, SATé outperforms all other methods in topological accuracy.

The improvement in topological accuracy using Canopy is more significant compared to native PRANK than compared to native PAGAN (Fig. 3a,b, Fig. S2). While this demonstrates that PAGAN’s modeling of alignment gaps is less sensitive to errors in the guide tree, this is at the expense of accuracy given a correct guide tree (data not shown).

**Figure 3:**
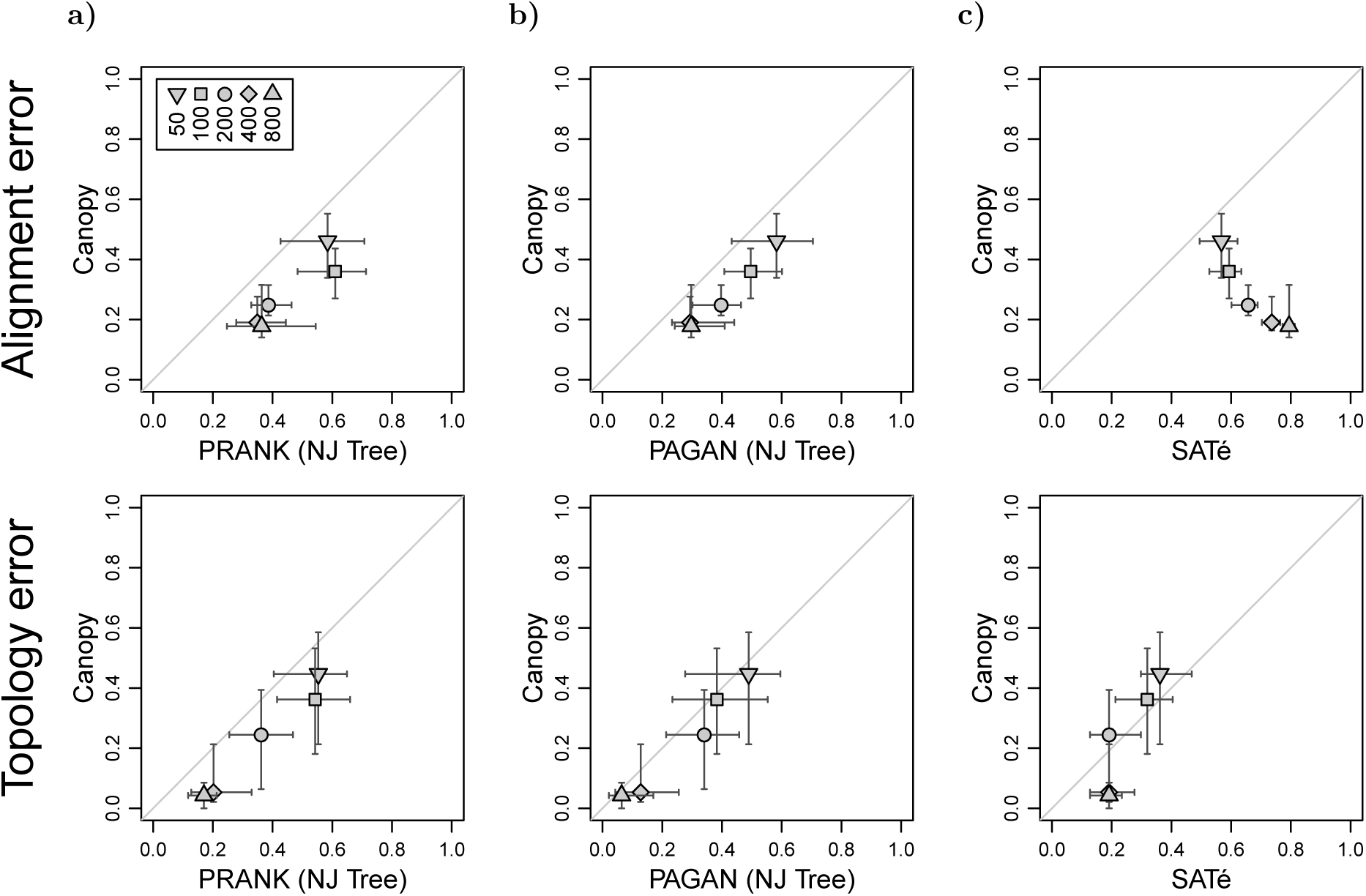
Relative performance of Canopy compared with (a) PRANK, (b) PAGAN and (c) SATé for ‘Intermediate’ data sets. The symbols indicate the median values and the errorbars show the second and fourth quartiles.

Our analyses with Canopy were based on 25 iterations, outputting the alignment-tree pair with the highest ML score. In line with Liu et al. (2012) we observed that the ML score correlates poorly with the accuracy of the alignment and the tree topology and a similar performance would have been observed by just accepting the solution after a few rounds of iteration. This is discussed with more detail in Supplemental Material.

## 3 Conclusions

Canopy provides a parallelized version of phylogeny-aware alignment with PRANK. Although the approach adds some overhead and Canopy’s performance does not scale linearly with the number of CPUs used, the tool provides a significant improvement over PRANK in the analysis of very large alignment data sets on modern computer hardware.

The phylogeny-aware algorithm relies strongly on the correctness of an alignment guide tree and, despite PRANK’s good performance in comparative analyses, it has been difficult to recommend the program for phylogenetic analyses. Indeed, we have shown this to be of genuine concern: when the guide tree is reflective of the true phylogeny, PRANK produces excellent alignments for inferring phylogenies, the topological correctness of which is comparable to trees inferred from true alignments. On the other hand, if the guide tree is erroneous, the alignment generated with PRANK may still be relatively accurate, but can contain errors that can seriously bias phylogenetic inference. One of the main reasons to reimplement the phylogeny-aware algorithm with PAGAN was to reduce sensitivity to errors in the guide tree. Our analyses demonstrate that this has largely been achieved by both PAGAN and Canopy for densely sampled sequences.

Canopy does not resolve all of the problems with sequence alignment, but it sets phylogeny-aware alignment methods on par with the leading alternative approach, SATé, enabling their use in phylogenetic analyses. In terms of topological accuracy, Canopy is superior to SATé in analysis of closely-related sequences: this level of divergence is achieved either by having low overall sequence divergence or very dense sampling of sequences. It has already been shown that the performance of phylogeny-aware alignment methods can degrade when the pairwise divergence of sequences grows and the insertion-deletion events cannot be correctly resolved (Löytynoja 2014). As our results show, this limitation can be largely resolved by including additional related sequences in the analysis.

Although Canopy is a significant improvement over PRANK with a guide tree inferred from the data, it is nowhere near the performance of using the true tree, so there is much room for improvement. As reported earlier (Liu et al. 2012), the correlation between the ML score used as an optimization criteria and the two accuracy measures is poor. Furthermore, without additional randomness added to the search, it is likely that all such approaches will get stuck in local optima and it is unclear whether a purely iterative approach is capable of resolving these issues.

It was recently suggested that the guide tree has no major impact on alignment accuracy and that a simple, comb-like guide tree would in fact produce the most accurate alignments (Boyce et al. 2014). While this may be permissible for some analysis problems, for example structural matching of protein sequences, our analyses support those of Tan et al. (2015) and clearly demonstrate that the effect of the guide tree on alignments used for phylogenetic inference is substantial. The phylogeny-aware algorithm relies on the correctness of the guide tree by design and is therefore particularly sensitive to guide tree error. However, when the guide tree is accurate and sequence sampling dense, the algorithm produces exceptionally good alignments for phylogenetic inference. Guide tree estimation, therefore, remains an important research question.

## 4 Acknowledgements

This work was supported by the Biocenter Finland, Biocentrum Helsinki and Marie Curie Career Integration Grants. We acknowledge CSC – IT Center for Science Ltd. for the allocation of computational resources.

